# Fear of missing out (FOMO) associates with reduced cortical thickness in core regions of the posterior default mode network and higher levels of problematic smartphone and social media use

**DOI:** 10.1101/2022.10.24.513508

**Authors:** Lan Wang, Xinqi Zhou, Xinwei Song, Xianyang Gan, Ran Zhang, Xiqin Liu, Ting Xu, Guojuan Jiao, Stefania Ferraro, Mercy Chepngetich Bore, Fangwen Yu, Weihua Zhao, Christian Montag, Benjamin Becker

## Abstract

Fear of missing out (FOMO) promotes the desire or urge to stay continuously connected with a social reference group and updated on their activities, which may result in escalating and potentially addictive smartphone and social media use. The present study aimed to determine whether the neurobiological basis of FOMO encompasses core regions of the reward or social brain, and the associations with the level of problematic smartphone or social media use. We capitalized on a dimensional neuroimaging approach to examine cortical thickness and subcortical volume associations in a comparably large sample of healthy young individuals (n = 167). Meta-analytic network and behavioral decoding were employed to further characterize the identified regions. Higher levels of FOMO associated with lower cortical thickness in the precuneus. In contrast, no associations between FOMO and variations in striatal morphology were observed. Meta-analysis decoding revealed that the identified precuneus region exhibited a strong functional interaction with the default mode network (DMN) engaged in social cognitive and self-referential domains. Together the present findings suggest that individual variations in FOMO are associated with the brain structural architecture of the right precuneus, a core hub within a large-scale functional network resembling the DMN involved in social and self-referential processes. FOMO may promote escalating social media and smartphone use via social and self-referential processes rather than reward-related processes per se.

## Introduction

Fear of missing out (FOMO) refers to the apprehension that others may have rewarding experiences from which oneself is absent (Przybylski, Murayama, DeHaan, & Gladwell, 2013, p. 1841). This apprehension in turn promotes the desire or urge to stay continuously connected with a social reference group and updated on their activities (Przybylski et al., 2013; Elhai, Yang, & Montag, 2021). Research on FOMO – often conceptualized as a rather stable trait-like characteristic (Wegmann, Oberst, Stodt, & Brand, 2017) – has rocketed in the context of growing concerns about excessive and potentially addictive social media and smartphone use (reviews see e.g. Elhai et al., 2021; Akbari et al., 2021). Accumulating evidence from different lines of research suggests an association between dispositional FOMO and problematic Internet use - in particular in the domains of social media and smartphone use - as well as associated emotional dysregulations and detrimental effects on daily life (Röttinger et al., 2021; Akbari et al., 2021; Rozgonjuk, Sindermann, Elhai, & Montag, 2020; Elhai, Yang, Rozgonjuk, & Montag, 2020; Yuan, Elhai, & Hall, 2021). Current conceptualizations propose that FOMO represents a general trait-like predisposition rendering individuals at an increased risk for excessive and problematic social media engagement and ultimately the development and maintenance of Internet Use Disorders (Akbari et al., 2021; Elhai et al., 2021; Röttinger et al., 2021). For associations between FOMO and age, gender and the Big Five of Personality see (Rozgonjuk, Sindermann, Elhai, & Montag, 2021)

From a conceptual perspective FOMO integrates two key motivational factors that may drive excessive and problematic social media use, specifically (1) reward-related components which refer to the fear of missing rewarding experiences, and (2) social-related components which refer to a strong need to belong to and stay connected with a social reference group (of note interactions with humans can have also a rewarding character). Whereas reward and motivation have traditionally received strong empirical attention as factors contributing to the development and maintenance of excessive and ultimately addictive engagement in behaviors and substance use (e.g. Brand et al., 2019; Koob & Volkow, 2016; Clark, Boileau, & Zack, 2018), social factors have only recently received increasing attention (e.g. Zhao et al., 2019; Zimmermann et al., 2019; Montag et al., 2018; Caouette & Feldstein Ewing, 2017; Gilman et al., 2016; Bedi et al., 2019).

On the neurobiological level the two domains and associated dysregulations have been mapped onto – at least to a certain degree – separable brain systems. Reward- and motivation-related processes have been strongly associated with the structure and function of the meso-cortico-limbic system encompassing the ventral tegmental area (VTA), striatum and orbitofrontal cortex (e.g. Koob & Volkow, 2016). Reward and motivational tasks reliably engage these circuits (e.g. Jauhar et al., 2021; Yang et al., 2021) and individual variations in reward-related processes have been associated in brain structural variations in these regions (e.g. Pehlivanova et al., 2018; Patel, Miles, & Nikolova, 2020). In the context of escalating use and addictive behaviors, studies have demonstrated that structural and functional variations in this circuitry predict escalating drug use (Becker et al., 2015; Bart et al., 2021) are manifest already during recreational engagement (e.g. X. Zhou et al., 2019; X. Zhou et al., 2020; F. Zhou et al., 2019) and represent a transdiagnostic marker across addictive disorders, including substance use disorders and behavioral addictions (Klugah-Brown et al., 2020; Solly, Hook, Grant, Cortese, & Chamberlain, 2021; Taebi et al., 2022; however see also (Klugah-Brown et al., 2021).

In contrast to the traditional emphasis on reward and motivational dysregulations in excessive and addictive behaviors, accumulating evidence emphasizes a contribution of social processes and social factors – in particular in adolescents and young adults – including the influence of peers and social stress, loneliness or the desire to gain social belongingness and status (e.g. Pokhrel, Fagan, Kawamoto, Okamoto, & Herzog, 2020; Snodgrass et al., 2018; Caouette & Feldstein Ewing, 2017). In support of this perspective FOMO has been associated with higher sensitivity to social exclusion (ostracism) as assessed with the cyberball-task (Holte, Fisher, & Ferraro, 2022). Although the underlying neurobiological systems overlap in some domains with those involved in reward processing (e.g. Martins et al., 2021) social processes such as mentalizing, self-reference, emotion recognition and social interaction have been robustly associated with distinct (social) brain systems, encompassing limbic, medial prefrontal and temporal regions as well as the large-scale default mode network (DMN) (e.g. Mihov et al., 2013; Amft et al., 2015; F. Zhou et al., 2020; Feng et al., 2021; Gan et al., 2022; Merritt, MacCormack, Stein, Lindquist, & Muscatell, 2021). Initial studies and conceptual reviews have begun to describe the neurobiological basis of dysfunctions in these domains in individuals with problematic substance use and reported for instance altered dorsomedial and limbic activation during social exclusion (e.g. Maurage et al., 2012), altered DMN recruitment during processing of social and self-referential information (e.g. Charlet et al., 2014; Hulvershorn et al., 2013; Zhao et al., 2019; overview see also Zhang & Volkow, 2019) or associations between empathic deficits and decreased brain volume in regions engaged in social cognitive processes (Baez et al., 2021).

In the context of the growing importance of FOMO in escalating and potentially addictive internet, in particular smartphone and social media use, the present study aimed to determine (1) the neurobiological basis of FOMO on the level of brain structure, (2) whether the neurobiological basis encompasses core regions of the reward or social brain, and (3) whether the neurobiological associations critically mediate the assumed association between FOMO and problematic smartphone or social media use, respectively. To this end we capitalized on a dimensional neuroimaging approach in a comparably large sample of healthy young individuals (n = 167, 72 females). The dimensional approach allows to examine associations between varying degrees in pathology related traits and symptoms and neurofunctional and neurostructural brain alterations (e.g. alexithymia or neuroticism, Li et al., 2018; Liu et al., 2021; Terock et al., 2020). Recently the approach has been increasingly applied and validated to determine brain structural and functional underpinnings of Internet and Social Media Addiction across the entire symptom range (Dong et al., 2021; Montag et al., 2018; X. Zhou et al., 2020). In line with the aims of the present study, our participants underwent a high-resolution brain structural MRI assessment, filled in inventories to assesses individual differences in levels of FOMO, and also smartphone and social media addiction were assessed using validated self-report scales. Our primary analyses examined associations between FOMO with cortical thickness and subcortical (striatal) brain volume and whether individual variation in these brain structural indices mediate the association between higher FOMO and excessive smartphone / social media use. Meta-analytic decoding was applied to provide a data-driven behavioral and functional characterization of the identified regions.

## Methods

### Participants and procedures

A cohort of 360 participants were selected from the database of the neuSCAN Lab at University of Electronic Science and Technology of China (UESTC). All individuals were healthy participants that had undergone structural MRI acquisition in the eight months (4.45±2.52) before the present study and were recontacted within the context of a larger project on the associations between FOMO and brain structure (the present research question) and a replication project for one of our previous studies on the association internet gaming tendencies and brain structure (X. Zhou et al., 2020; replication in Klugah-Brown et al., in preparation). After initial quality assessment of the brain anatomical images and excluding duplicates 287 participants were contacted via phone and invited to participate in an online self-report assessment including questionnaires, data from 204 receipts were collected. Finally, 167 eligible participants (mean age=21.35±3.17, 72 females) in total were included in subsequent data analyses for the current study after removal of 37 invalid questionnaires. Inclusion criteria included the absence of a history of psychiatric disorders and the usage of psychotropic substances (**Figure 1**.). The protocols had ethical approval by the University of Electronic Science and Technology of China and were in accordance with the latest Revision of the Declaration of Helsinki.

**Figure 1.**
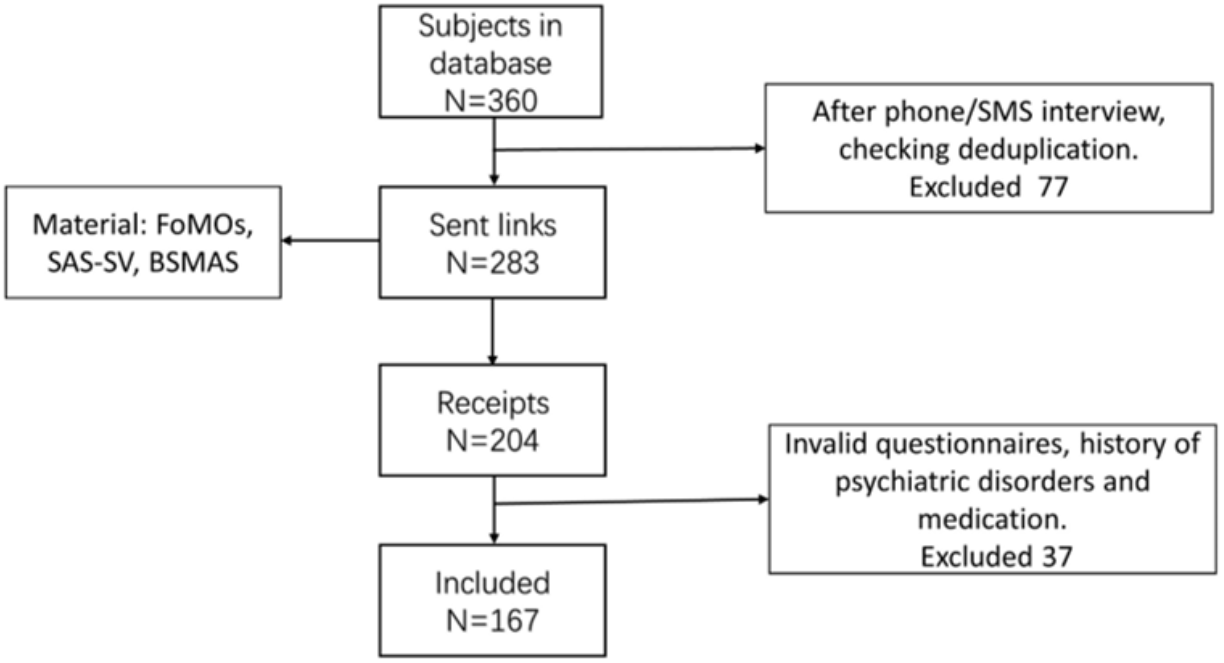
Exclusions and procedures for collecting questionnaire data

### Assessment of FOMO and related variables

In line with our study aims participants were administered the Fear of Missing Out scale (FOMOs), Smartphone Addiction Scale-Short Version (SAS-SV), Bergen Social Media Addiction (BSMAS) and demographic questions, including sex, age, ethnicity, and education level.

Individual variations in the trait-like characteristic FOMO (Akbari et al., 2021; Elhai et al., 2021) were assessed using the validated FOMOs developed by Przybylski et al., which has a demonstrated high sensitivity and reliability (Przybylski et al. 2013). The scale presents 10 items scored on a five-point Likert-type scale ranging from ‘Not at all true of me’ to ‘Extremely true of me’. We used the validated Chinese version of the SAS-SV for the present study population (Xie, Wang, Wang, Zhao, & Lei, 2018). Problematic smartphone sue was assessed using the SAS-SV, a short version of the 33 items Smartphone Addiction Scale (Kwon et al., 2013). While the SAS-SV only retains 10 items it incorporates all core factors from the original version (i.e. daily-life disturbance, positive anticipation, withdrawal, cyberspace-oriented relationship, overuse, and tolerance) (Choi, Kwon, Kim, Cho, & Yang, 2013) assessed via a six-point Likert scale ranging from 1 (strongly disagree) to 6 (strongly agree). The SAS-SV has been validated in various languages and has shown a high reliability and validity. We used the Chinese version, translated and validated preciously (B. Chen et al., 2017).

The BSMAS is an adapted version of the Bergen Facebook Addiction Scale (BFAS) (Andreassen, Torsheim, Brunborg, & Pallesen, 2012), replacing the keyword ‘Facebook’ with ‘social media’ (Andreassen, Pallesen, & Griffiths, 2017). BSMAS contains 6 items that correspond to 6 essential addiction symptoms namely salience, conflict, mood modification, withdrawal, tolerance, and relapse in the context of using social media use during the last year assessed via a 5-point Likert scale ranging from 1 (very rarely) to 5 (very often). The Chinese version of BSMAS was utilized in the current study (I. H. Chen et al., 2020). Internal consistencies are reported in Table 2.

### Structural MRI data acquisition and preprocessing

All participants underwent high-resolution brain structural assessments on a 3.0 Tesla GE MR750 system (General Electric Medical Systems, Milwaukee, WI, USA). T1-weighted images of the whole brain were acquired using the following acquisition parameters: voxel size=1.5mm^3^, repetition time (TR)=2300ms, echo time (TE)=2000ms, flip angle=9 degrees, field of view (FOV)=256×256mm, acquisition matrix=256×256, slice thickness=1mm, number of slices=156. Structural MRI data were preprocessed with CAT12, a computational anatomy toolbox (https://neuro-jena.github.io/cat) for structural brain images based on Statistical Parametric Mapping software (SPM12, Welcome Department of Cognitive Neurology, London, UK, https://www.fil.ion.ucl.ac.uk/spm/software/spm12). CAT12 provides a fully automated method to estimate cortical thickness and the central surface of hemispheres based on the projection-based thickness method. The preprocessing was implemented according to recommendations in the CAT12 surface-based morphometry (SBM) preprocessing pipeline manual, including cortical surface estimation, topological correction, and spherical mapping and registration. Cortical thickness was smoothed using 15mm full width at half maximum (FWHM) Gaussian kernel (Dahnke, Yotter, & Gaser, 2013). Given previous reports on associations between problematic internet engagement and striatal volume (e.g. X. Zhou et al., 2020) we additionally assessed mean tissue volumes in subcortical regions in line with the standard procedures in the CAT12 manual (see also (Gaser & Dahnke, 2016), including segmentation into gray matter (GM), white matter (WM), and cerebrospinal fluid (CSF). GM volumes from the striatum as a single region of interest (ROI) were extracted in native space. For further validation of associations with striatal subregions we separately extracted the volumes of the dorsal and ventral striatum from the modulated normalized GM segments in standard space using the ROI Signal Extractor tool of Data Processing and Analysis for Brain Imaging (DPABI, http://rfmri.org/DPABI; (Yan, Wang, Zuo, & Zang, 2016). Normalization and preprocessing for this analysis were conducted in line with recommendations in the CAT manual (details of the processing pipeline see also (X. Zhou et al., 2022). The dorsal and ventral striatum subregions were defined based on the Brainnetome atlas (Fan et al., 2016).

### Statistical analyses

Multiple linear regression models were performed in SPM12. Associations between FOMO and cortical thickness at the whole-brain level were first explored controlling for sex, age, and image quality rating (IQR) as covariates. Next, SAS-SV and BSMAS were included as covariates to validate the robustness and specificity of the associations between FOMO and cortical thickness. Results were thresholded at a cluster-level p<0.05 with family-wise error (FWE) correction (initial voxel-level threshold at p<0.001). Our a priori hypothesis on associations between FOMO and the volume of the striatum as subcortical core reward processing region was examined using a Pearson correlation analysis in SPSS 21. Next, mediation analyses were conducted using the PROCESS tool implemented by SPSS, and the 95% confidence interval (Hayes) including non-zero values is considered as significant results (Hayes, 2012).

### Meta-analytic network and behavioral decoding of the identified region

To quantitatively characterized the functional networks and mental processes associated with the regions displaying FOMO associations we conducted meta-analytic decoding across imaging studies from the literature with the Neurosynth platform (http://www.neurosynth.org; Yarkoni, Poldrack, Nichols, Van Essen, & Wager, 2011). The identified peak coordinate was subjected to the meta-analytic module to generate meta-analytic maps of other brain regions coactivated with the defined area, including functional connectivity and coactivation maps. After combination of these two maps the results were visualized with the Anatomy3.0 toolbox (Eickhoff et al., 2005) as implemented in SPM12. We next employed meta-analytic decoding using the “Decoder” module to characterize the associated behavioral functions.

### Ethics

The study and its procedures had full approval by the local ethics committee and adhered to the most recent version of the Declaration of Helsinki and all participants were required to provide informed consent.

## Results

### Demographics and descriptive statistics of questionnaires

N = 167 subjects were included, n=95 male, descriptive information of each questionnaire see **Table 1**, including FOMO (29.76±6.94), smartphone addiction (34.76±8.48), and social media addiction (15.61±4.51). No sex differences were observed for FOMO (t = -0.028, p = 0.978), smartphone (t = 0.234, p = 0.815) and social media addiction (t = -0.623, p = 0.815). Scales in the present study had good reliability and validity, and there were high positive correlations between SAS-SV and FOMO, BSMAS and FOMO (**Table 2**. Spearman correlations remained stable considering non-normal distributions). implying that supplementary controls for the SAS-SV and BSMAS is crucial in the subsequent analyses to control for common or indirect effects of these constructs on brain volume.

**Table 1.** Demographic characteristics of participants.

|  | N | Age | FOMOs | SAS-SV | BSMAS |
| --- | --- | --- | --- | --- | --- |
|  | 167 | 21.35±3.17 | 29.76±6.94 | 34.76±8.48 | 15.61±4.51 |
| **Male** | 95 | 21.44±2.85 | 29.75±7.13 | 34.89±8.29 | 15.42±4.72 |
| **Female** | 72 | 21.22±3.57 | 29.78±6.73 | 34.58±8.77 | 15.86±4.25 |

**Table 2.** Reliability, validity and correlation matrix of questionnaires.

|  | Cronbach's |  |  | FOMOs | SAS-SV | BSMAS |
| --- | --- | --- | --- | --- | --- | --- |
|  | Alpha | KMO |  |  |  |  |
| **FOMOs** | 0.791 | 0.782 |  | -- |  |  |
| **SAS-SV** | 0.826 | 0.834 | **Correlation** | 0.440** | -- |  |
| **BSMAS** | 0.817 | 0.815 |  | 0.246** | 0.579** | -- |

*Note:* \*\* Correlation is significant at the 0.01 level (2-tailed).

### Negative association between cortical thickness and FOMO

A significant negative association between FOMO and cortical thickness in the right precuneus was found controlling for age, sex, IQR as covariates (peak voxel t=4.40, cluster-level pFWE<0.05, **Table 3**). Notably, the association remained stable when smartphone addiction and social media addiction were additionally controlled as covariates (peak voxel t=4.30, cluster-level pFWE<0.05). Despite the high correlation between FOMO and excessive smartphone or social media use on the behavioral level a mediation analysis did not reveal a significant mediation effect of precuneus thickness on this association (95% CI -0.058 to 0.003 for SAS-SV, 95% CI -0.115 to 0.016 for BSMAS). Together this indicates that higher FOMO is – to a certain extent – specifically associated with thinner cortical thickness of the right precuneus (**Figure 2**).

**Table 3.**
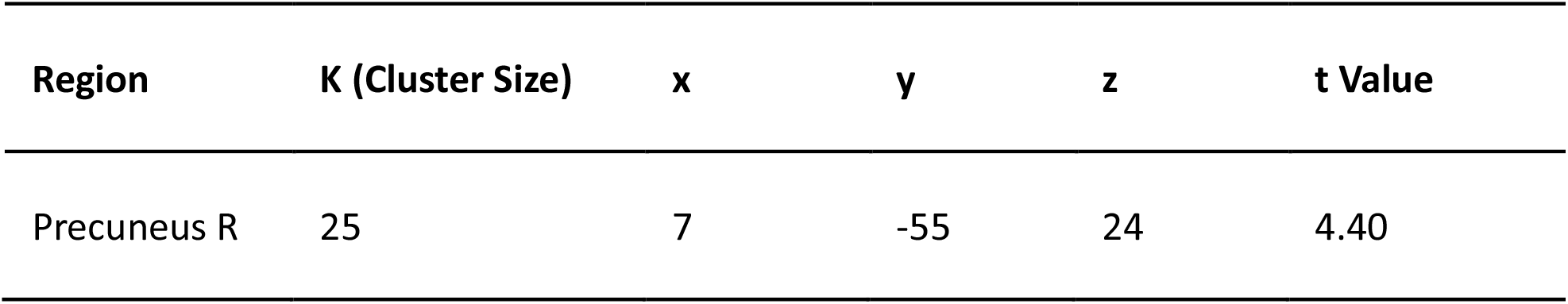
Significant association between FOMO and cortical thickness of the right.

**Figure 2.**
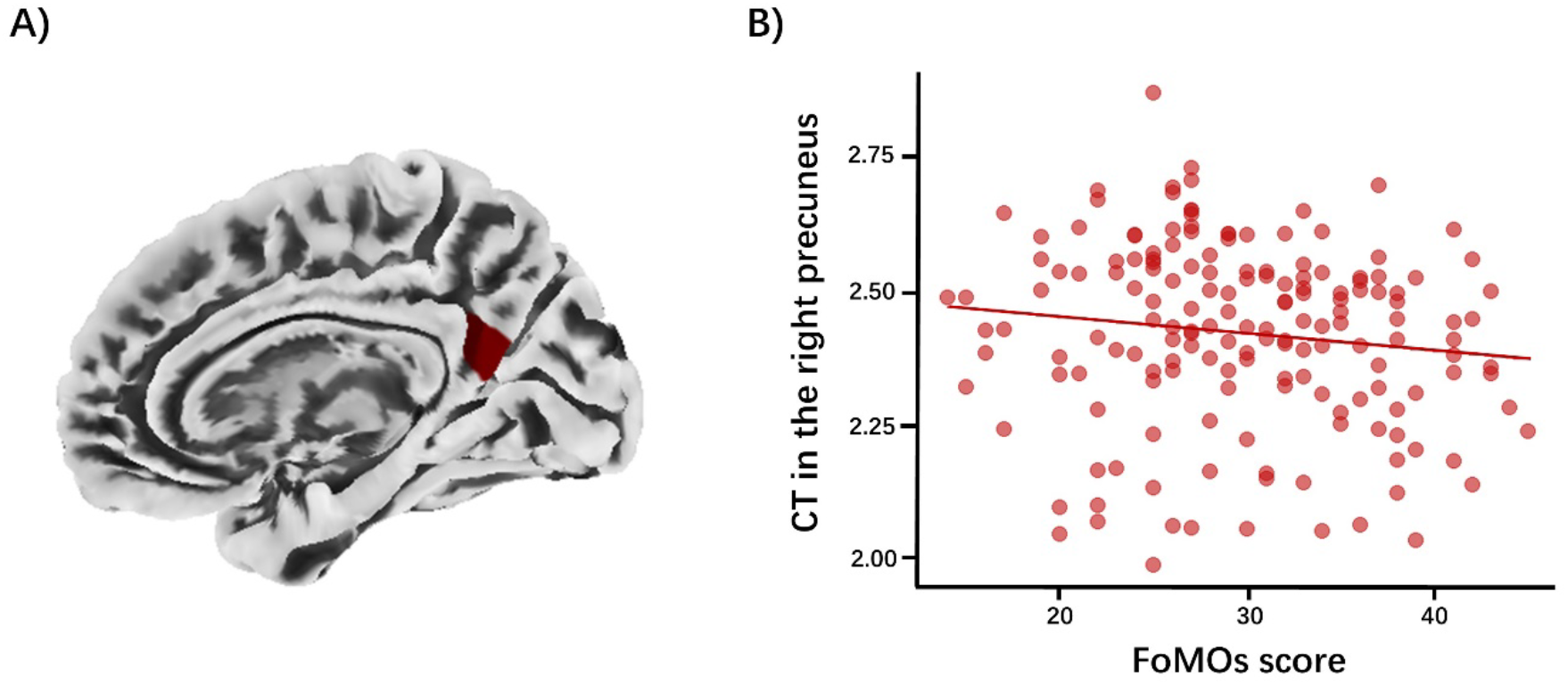
Association between higher FOMO and thinner cortical thickness of the precuneus **A)** Depicts the negative association between FOMO cortical thickness in the right precuneus at pFWE<0.05 corrected threshold level. **B)** Presents a scatter plot between extracted surface thickness measures and the individual FOMO scores for visualization purpose.

### No significant association between FOMO and striatal volume

There was no significant correlation between extracted striatal gray matter volume and levels of FOMO (r=0.073, p=0.346; r=0.04, p=0.660 after additionally controlling for age and sex as covariates), arguing against strong morphological associations between FOMO and volume of the striatal system. Additional control analyses did furthermore not reveal associations with the excessive smartphone or social media use as assessed by the SAS-SV or the BSMAS, respectively (all ps > 0.364). Furthermore, the results remained stable in additional control analyses that examined extracted gray matter volumes for the dorsal and ventral striatum separately (see **Table 4**).

**Table 4.** Correlation between FOMO and the striatum.

|  | Striatum | Dorsal striatum | Ventral striatum |
| --- | --- | --- | --- |
| **FOMO** | r=0.073 | r=0.067 | r=0.109 |
|  | p=0.346 | p=0.392 | p=0.161 |

### Meta-analytic decoding: network level and behavioral characterization of the right precuneus

Computing the conjunction between the connectivity and co-activation meta-analytic characterization maps for the identified right precuneus region revealed that this region exhibited strong and spatially specific functional interactions with core regions of the default mode network (DMN) (**Figure 3A))**. The identified network included key nodes of the core DMN, including the precuneus/posterior cingulate cortex (PCC), medial prefrontal cortex (mPFC) and angular gyrus as well as the dorsal medial DMN subsystems such as the dorsal medial prefrontal cortex (dmPFC) and the medial temporal DMN such as the hippocampus. Behavioral decoding of this network correspondingly revealed DMN-associated functions, including social cognitive functions such as theory of mind, social, mentalizing and self-referential functions such as autobiographic memory or self-referential (**Figure 3B))**.

**Figure 3.**
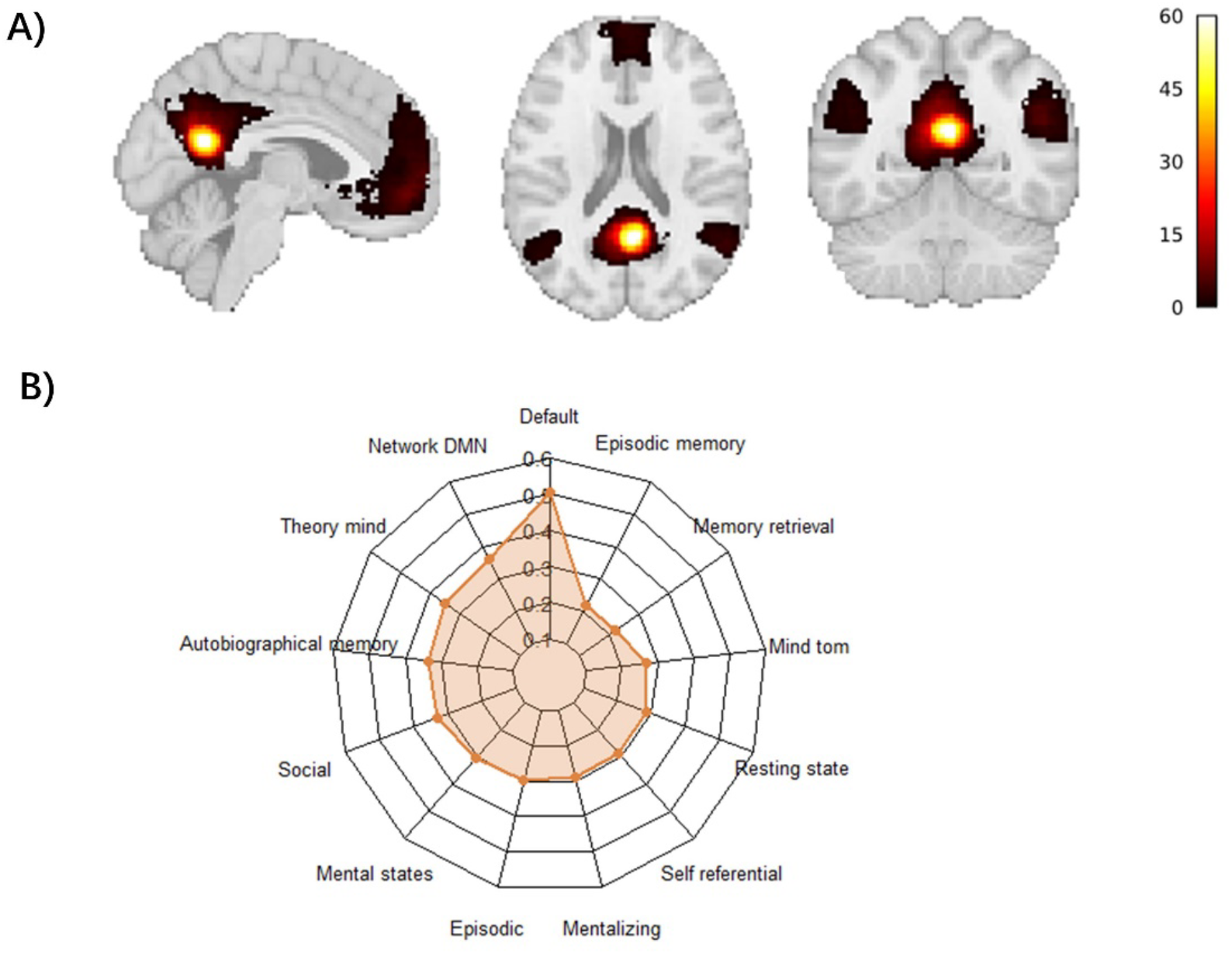
Network-level and behavioral decoding of the identified right precuneus region **A)** Overlap of the functional connectivity and co-activation maps of the precuneus identified the key regions of the default mode network. **B)** Radar chart illustrating the meta-analytically derived functional characterization of the network interacting with the right precuneus (13 terms were retained from 20 after excluding terms repetitions from the meta-analytical decoding).

## Discussion

Utilizing a dimensional neuroimaging approach in a comparably large sample the present study allowed – to our knowledge - for the first time to determine a brain structural correlate of FOMO. FOMO has received increasing interest within the context of growing concerns about problematic and escalating social media and smartphone usage among young individuals (Elhai et al., 2021; Przybylski et al., 2013). In line with our hypothesis higher levels of FOMO were associated with higher levels of both, problematic smartphone, and social media usage. On the brain structural level individual variations in FOMO varied as a function of regional-specific variations in cortical thickness, such that higher levels of FOMO associated with lower thickness of the right precuneus. Despite the association on the behavioral level controlling for problematic smartphone or social media use did not change the neural results and cortical thickness of the precuneus did not mediate the association between FOMO and smartphone or social media usage. Meta-analytic network and behavioral decoding revealed that the identified precuneus region exhibited a strong functional interaction with the default mode network (DMN) characterized by a robust involvement in a range of social cognitive and self-referential behavioral domains. In contrast, no associations between FOMO and variations striatal volume were observed. Together the present findings suggest that individual variations in FOMO are associated with individual variations in the brain structural architecture of the right precuneus, a core region in strong functional interaction with a large-scale functional network resembling the DMN and involved in social and self-referential processes.

In line with a growing number of studies we observed a positive association between higher levels of FOMO and more problematic smartphone and social media usage. On the one hand, FOMO has for instance been associated with development of problematic usage of a number of internet applications, including problematic internet usage (Röttinger et al., 2021) and smartphone addiction (Alinejad, Parizad, Yarmohammadi, & Radfar, 2022). However, on the other hand, an increasing number of studies underscore the role of social and self-related cognitions in this association such that the association between FOMO and smartphone addiction is mediated by loneliness (Alinejad et al., 2022) while FOMO mediates the association between parental support, self-construal, self-concept clarity or interpersonal sensitivity with problematic smartphone use (Kim, 2022; Lin et al., 2021; Servidio, Sinatra, Griffiths, & Monacis, 2021).

On the neural level we demonstrate for the first time an association between higher levels of FOMO and regional-specific lower cortical thickness of the right precuneus. The precuneus represents a functionally heterogenous region that has been involved in an entire range of highly integrative domains, ranging from basic cognitive functions such as attention towards complex social functions, including mentalizing and self-referential processes (e.g. Menon, Rivera, White, Glover, & Reiss, 2000; Schurz, Radua, Aichhorn, Richlan, & Perner, 2014; Zhao et al., 2019). From a system level perspective, the precuneus forms – together with the posterior cingulate cortex –represents the integral functional hub of the posterior DMN(Andrews-Hanna, Smallwood, & Spreng, 2014). The DMN encompasses a set of medial frontoparietal networks, including key functional hubs located in the precuneus/posterior cingulate cortex, the angular gyrus and the medial prefrontal cortex as well as a dorsal medial subsystem encompassing regions such as the dorsal prefrontal cortex and a medial temporal subsystem encompassing regions such as the hippocampus which closely resemble the network identified in our meta-analytic network characterization of the right precuneus. The DMN plays a key role in internally directed and self-generated thoughts and has been involved in an entire range of social and self-referential domains, including social interaction, social exclusion, social isolation and loneliness (e.g. Feng et al., 2021; Morr, Liu, Hurlemann, Becker, & Scheele, 2022; Mwilambwe-Tshilobo & Spreng, 2021; Spreng et al., 2020). The core system including the precuneus / posterior cingulate node has been proposed to support in particular self-referential processes, mentalizing and autobiographic and social information processing (Andrews-Hanna et al., 2014). Together with the highly convergent behavioral characterization of the identified network in the present study these results may reflect that FOMO associates with variations in the neurobiology of a core hub underlying self-referential and social processing.

In contrast to our hypothesis, we did not find associations between FOMO and striatal volume which have been frequently reported in individuals with problematic behavior, including problematic or addictive internet use, gaming and social-media engagement (e.g. Yu et al., 2022; X. Zhou et al., 2020; X. Zhou et al., 2019), which may suggest that FOMO has a differential neurobiological basis compared to addictive tendencies. This is further underscored by the observation that controlling for social media and smartphone use disorders did not change the association between FOMO and precuneus thickness. This also speaks for the fact that FOMO is a psychological construct, which can be seen independent of online use and of course also plays a pivotal role in every day life, when interacting with humans. Of note, disentangling trait and state FOMO as put forward by (Wegmann et al., 2017) would be highly interesting in a further study. In the updated FOMO scale by Wegmann et al. trait FOMO refers to general fear of missing out not necessarily linked to FOMO in the online world, whereas additional formulated items in the state FOMO scale aim to assess FOMO in the online realm. It will be interesting to see if trait/state FOMO as operationalized in the Wegmann et al. (2017) work will provide additional insights into the underlying biology of FOMO. Finally, from a conceptual level the present findings may suggested that FOMO may exert its potential influence on social media and smartphone use rather via social domains than reward-related dysfunctions which are commonly considered as a transdiagnostic vulnerability for addictive behavior.

The findings of the present study need to be considered in the context of limitations. Although the present study encompassed a comparably large sample size the findings require an independent replication. Behavioral measures assessing social functioning, self-referential processing or reward sensitivity were not assessed and thus future studies need to employ a more careful behavioral characterization of FOMO.

## Authors’ contribution

LW, XZ, BB, and CM conceptualized, designed, and wrote the initial protocol and draft. LW, XZ conducted and validated the formal analysis. XS, XG, RZ, XL, TX, GJ, FY, SF, MCB and WZ carried out the experiments, MRI acquisition and provided technical support. WZ, BB, and CM reviewed and revised the manuscript. BB obtained funding and administrated the project. All authors contributed to and approved the final version of the manuscript, and take responsibility for the integrity of the data and the accuracy of the data analysis.

